# Pulsed ultrasound modulates the cytotoxic effect of cisplatin and doxorubicin on cultured human retinal pigment epithelium cells (ARPE-19)

**DOI:** 10.1101/2023.10.02.560520

**Authors:** Seyed Omid Mohammadi, Megan C. LaRocca, Christopher D. Yang, Jordan Jessen, M. Cristina Kenney, Ken Y. Lin

## Abstract

**Objective:** Pulsed ultrasound has been proposed as a tool to enhance ocular drug delivery, but its effects on drug potency are not well understood. Doxorubicin-HCl and cisplatin are two drugs commonly used to treat ocular melanoma. We report the effects of pulsed ultrasound on the cytotoxicity of doxorubicin-HCl and cisplatin *in vitro*.

**Methods:** Cultured human retinal pigment epithelium cells (ARPE-19) cells were treated with doxorubicin-HCl or cisplatin in the presence or absence of ultrasound. MTT and Trypan blue assays were performed at 24 and 48 hours post-treatment to assess cell metabolism and death.

**Results:** Cells treated with ultrasound plus doxorubicin-HCl demonstrated a significant decrease in metabolism compared to cells treated with doxorubicin-HCl alone. In contrast, cells treated with ultrasound plus cisplatin exhibited a significant increase in metabolism compared to cells treated with cisplatin alone at 48-hours. Cells treated with cisplatin pre-treated with ultrasound (US-Cis) exhibited a significant decrease in metabolism. Cell death was similar in doxorubicin- and cisplatin-treated cells with and without ultrasound.

**Conclusion:** Pulsed ultrasound enhances the cytotoxicity of doxorubicin-HCl at 24- and 48-hours post-treatment but abrogates cisplatin toxicity 48-hours post-treatment. This suggests ultrasound modulates cell-drug interactions in a drug-specific manner. These findings may influence the future development of ultrasound-assisted ocular drug delivery systems.

## Introduction

Since the first ocular echogram was performed in 1956, ultrasound has become a valuable tool for visualizing ocular structures. The anatomy of the eye facilitates insonation in several ways. Its superficial placement allows ultrasound waves to travel without obstruction, and the physical properties of its fluid-filled chambers allows facile ultrasound transmission to both the anterior and posterior segments. Moreover, ultrasound probes are mobile, which allows operators to approach the eye at different incident angles to capture high-quality images [1]. These features make ocular ultrasound an indispensable diagnostic tool.

Although ultrasound is most often used as an imaging modality, its therapeutic role in ophthalmology is not fully defined [2]. One recent development relevant to therapeutic ultrasound in the eye involves high-intensity focused ultrasound (HIFU) to treat glaucoma [3]. HIFU coagulates the ciliary body to modulate aqueous humor production and subsequently reduces intraocular pressure [3–5]. The use of ultrasound has also been studied in the context of ocular drug delivery. Methods of ultrasound-targeted drug delivery have included nano- and microparticles, including liposomes and micelles, phase-change agents, and thermally responsive materials [6]. Notably, ultrasound-sensitive monodisperse microbubbles have been noted as optimal for ultrasound-mediated drug delivery as their acoustic shadows allow ultrasound operators to selectively target tissues [7]. Due to the complex anatomy and physiology of the eye, drugs are often absorbed in subtherapeutic doses due to physical obstruction [8]. To achieve an adequate therapeutic window in ocular tissues, many compounds would need to be infused intravenously at such high concentrations as to cause significant systemic side effects. Ultrasound-assisted drug delivery in the eyes represents a potential solution to this challenge.

Few studies have assessed the relationship between pulsed ultrasound and antineoplastic agents *in vitro*. Prior studies have demonstrated that low-level ultrasound in combination with adriamycin and diaziquone reduces uterine cervical squamous cell tumor size reduction *in vitro* and *in vivo* [9]. Ultrasound has also been demonstrated to enhance the delivery of dexamethasone sodium phosphate through the cornea *in vivo* [10]. Yet, no reported studies have explored the impact of pulsed ultrasound on the cytotoxicity of antineoplastic drugs in cultured human retinal pigment epithelium (ARPE-19) cells. Due to its established cytotoxicity, doxorubicin, an anthracycline, was chosen as a model compound for our *in vitro* study. Cisplatin, a platinum analog and alkylating agent with known retinotoxic effects, including retinopathy, ischemia, and hemorrhage, was also evaluated [11]. These two drugs were chosen for our study because they have well-defined toxicity profiles and dose-response effects in the ARPE-19 cell line, and therefore serve as useful tools to interrogate the effect of pulsed ultrasound on their cytotoxicity.

The goal of the present study is to characterize the effects of pulsed ultrasound in the presence of doxorubicin and cisplatin as a method to modulate drug toxicity in ARPE-19 cells and expand upon its potential therapeutic role in ocular drug delivery.

## Materials and Methods

### Pulsed Ultrasound Apparatus

Our experimental ultrasound apparatus included a room-temperature water bath with a 0.5-inch inlet on its inferior surface for the ultrasound transducer to enter; a stand to elevate and stabilize the water bath and transducer, and a rig to fix the inverted cell plate at the focal point of the transducer (Figure 1). ARPE-19 cells [ATCC® CRL-2302™; American Type Culture Collection, Manassas VA] were plated on 96-well plates with 360 μL of media. Cell plates were saturated with culture media and sealed with MicroAmp^TM^ Optical Adhesive Film (Thermo Fisher Scientific, Austin, TX), inverted, and submerged in the water bath to mitigate potential noise from the air-liquid interface and maximize ultrasound penetrance. After inversion, cell plates were fixed 1 inch from the transducer and displaced laterally in a controlled fashion for 5 minutes at a constant velocity to ensure uniform ultrasound exposure (Figure 1). A portable Olympus EPOCH 650 system (Waltham, MA) was used to generate ultrasound waves. ARPE-19 cells were initially subject to ultrasound at a frequency of 1 MHz, power of 4.93 W/cm^2^, and pulse repetition frequency (PRF) of 100 Hz for 5 minutes based on prior ultrasound experiments on ARPE-19 cells reported in the existing literature [12]. Experiments were then repeated with the same parameters but at a PRF of 30 Hz because ultrasound treatment at a PRF of 100 Hz exerted a significant decrease in cell metabolism at baseline (Figure 2b). 3-[4,5-dimethylthiazol-2-yl]-2,5 diphenyltetrazolium bromide (MTT) assay was performed at 24 and 48 hours to assess the effects of ultrasound on cell metabolism.

**Figure 1:**
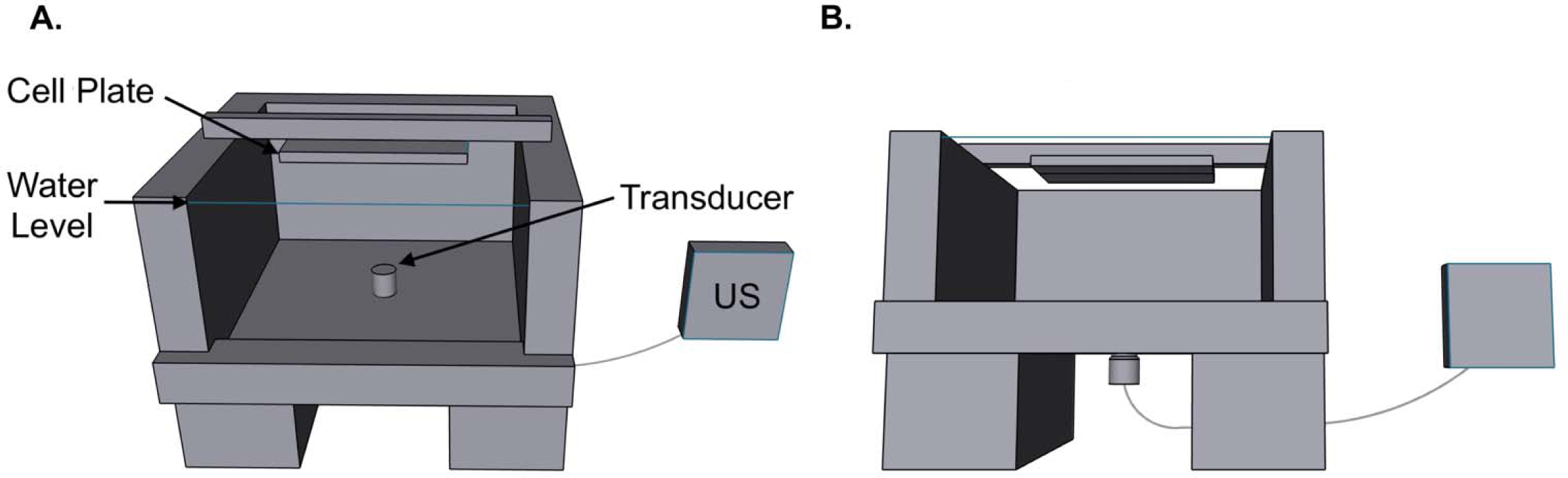
Three-dimensional CAD render of the experimental ultrasound (US) set-up. Inverted cell plates were placed 1 inch from the transducer tip at its focal point. The blue line denotes a water level allowing full submersion of the cell plate. The apparatus allowed only lateral movement (in the x- and y-direction) of the cell plate and maintained a fixed 1-inch distance between the cell plate and transducer.

**Figure 2:**
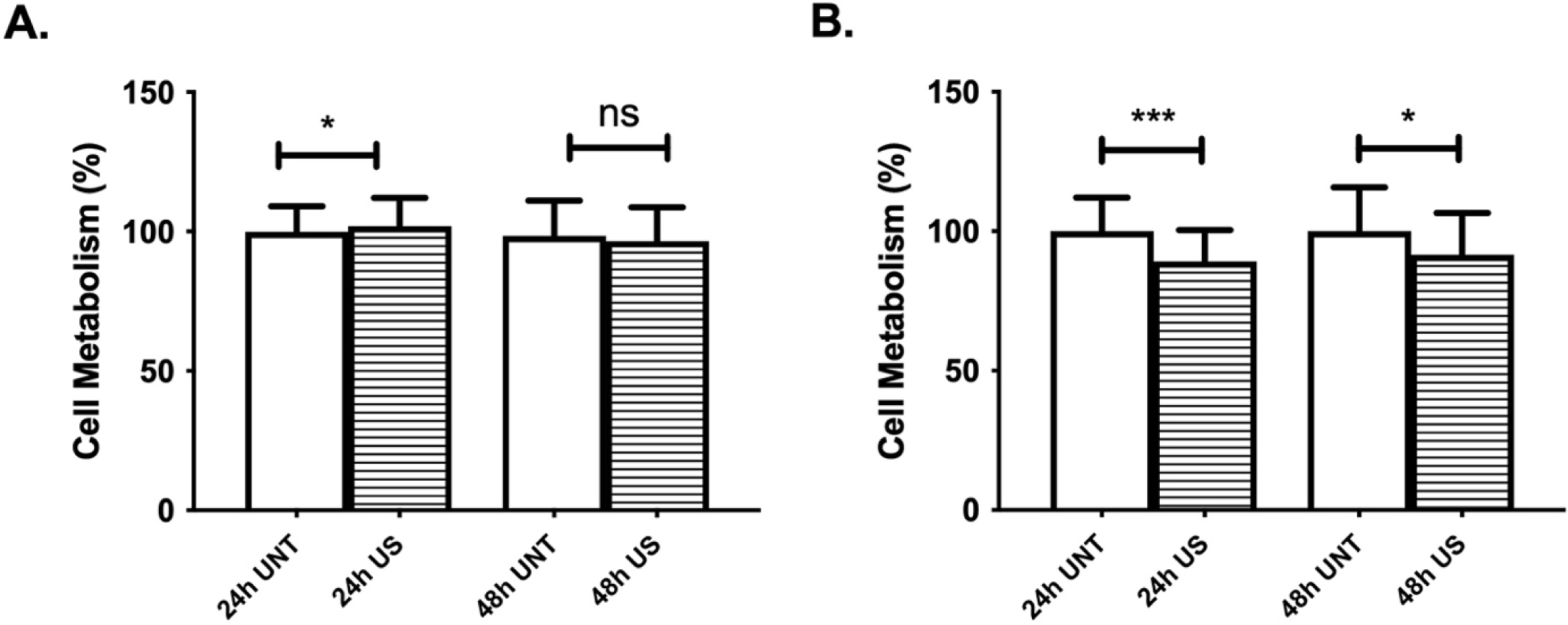
The effects of ultrasound treatment alone on ARPE-19 cell metabolism at a pulse frequency of 1 MHz, power of 4.93 W/cm^2^, and a PRF of 30 Hz (A) and 100 Hz (B). After 48 hours, there was no significant decrease in cell metabolism at a PRF of 30 Hz (p=0.1398). There was a significant decrease in cell metabolism at a PRF of 100 Hz (24 hours, p=0.0002; 48 hours, p=0.0181). These results confirmed the effectiveness of our ultrasound setup.

### Doxorubicin-HCl and Cisplatin Toxicology

Doxorubicin-HCl toxicity in ARPE-19 cells was evaluated before pulsed ultrasound experiments. 0.01, 0.1, 1, 10, and 100 μg/mL doxorubicin-HCl were chosen as working concentrations [13]. A stock doxorubicin-HCl solution was initially created by dissolving doxorubicin-HCl (Fisher Scientific, Hanover Park, IL, United States) in DMSO; this solution was serially diluted in media to yield the working concentrations listed above. ARPE-19 cells were plated in 96-well plates containing 360 μL doxorubicin-HCl solution or media. Untreated cells were plated with media alone as experimental controls. After a 24- or 48-hour incubation period, MTT assays were performed to establish the baseline toxicity of doxorubicin-HCl on ARPE-19 cells. All doxorubicin-HCl experiments were repeated three times. Cisplatin (BluePoint Laboratories, Little Island Cork, Ireland) was used at working concentrations of 20 μM and 40 μM [14].

### Doxorubicin-HCl, Cisplatin, and Pulsed Ultrasound Treatment

ARPE-19 cells were treated with doxorubicin-HCl or cisplatin with or without pulsed ultrasound exposure. ARPE-19 cells were initially plated in 96-well plates at a density of 1×10^5^ cells/mL and treated with 360 μL doxorubicin-HCl or cisplatin solution. Doxorubicin-HCl was used at the concentrations described in *Doxorubicin-HCl and Cisplatin Toxicology*. After loading wells with doxorubicin-HCl or cisplatin, cell plates were treated with 1 MHz ultrasound at a power of 4.93 W/cm^2^ and PRF of 30 Hz. Pulsed ultrasound was not applied to control wells. Media were removed and wells were refreshed with doxorubicin-HCl or cisplatin solution. Plates were then incubated for 24 or 48 hours at 37 degrees Celsius and 5% CO2 saturation. Following incubation, MTT assay was performed to assess metabolic activity as a proxy for cell viability. To determine if pulsed ultrasound treatment influences cisplatin toxicity, experiments were repeated utilizing pulsed ultrasound-treated cisplatin (US-Cis). First, cisplatin was treated with 1 MHz ultrasound at a power of 4.93 W/cm^2^ and PRF of 30 Hz. US-Cis was then added to the cultured cells. Plates were inverted and placed in the water bath, but no further pulsed ultrasound was applied to experimental groups. Plates were incubated for 48 hours, and cell metabolism was assessed via the MTT assay described below.

### Measurement of Cell Metabolism (MTT Assay)

The MTT (3-[4,5-dimethylthiazol-2-yl]-2,5 diphenyltetrazolium bromide) assay, based on the conversion of MTT to insoluble formazan crystals via NADPH-dependent oxidoreductases, was used to assess cellular metabolism as a proxy for cell viability as described by T. Mosmann [15]. MTT assay was primarily used to evaluate the *in vitro* cytotoxic effects of antineoplastic drugs. Assay outputs were quantified by a BioTek ELx808 spectrophotometer at 570 nm and 630 nm. Results were expressed as percent cell viability relative to untreated controls. Each experiment was replicated three times.

### Measurement of Cell Death (Trypan Blue Assay)

Trypan blue exclusion assays were conducted to assess cell death. 5×10^5^ ARPE-19 cells were plated in 6-well plates. After a 24-hour incubation period, media were replaced with either doxorubicin-HCl (100 μg/mL or 1 μg/mL) or cisplatin (40 μM or 20 μM). Half of the wells were treated with 90-seconds of 1 MHz ultrasound at a PRF of 30 Hz. The medium in each well was aspirated to a final volume of 2 mL.

Trypan blue exclusion assay, as described by Strober [16], was performed 24- and 48-hours post-treatment and read using a Vi-CELL XR Cell Viability Analyzer (Beckman Coulter).

### Statistical Analysis

Unpaired two-tailed tests were used for statistical analysis via Prism (GraphPad Software, San Diego, CA). P-values <0.05, <0.01, <0.001, <0.0001 were denoted as *, **, ***, and ****, respectively and all denote statistical significance; non-significance was denoted as “ns”. All values were expressed as average +/-standard deviation.

## Results

### Ultrasound Treatment

Our pulsed ultrasound apparatus established a sealed liquid-liquid interface for facile pulsed ultrasound application *in vitro*. At 48 hours post-treatment, pulsed ultrasound at a frequency of 1 MHz, power of 4.93 W/cm^2,^ and PRF of 30 Hz did not significantly change ARPE-19 cell metabolic activity (Figure 2a). At 24 hours post-treatment, pulsed ultrasound with a PRF of 30 Hz caused a significant increase in ARPE-19 cell metabolism compared to the untreated control (p=0.0340; Figure 2a). This observation confirms that pulsed ultrasound, when applied at the described parameters, does not decrease metabolic activity in ARPE-19 cells. Yet, when PRF was increased to 100 Hz, significant decreases in cell metabolism were observed via MTT assay at 24- and 48-hours post-treatment (24 hours, p=0.0002; 48 hours, p=0.0181, Figure 2b).

### Doxorubicin-HCl Toxicology

MTT assay revealed a generally dose-dependent decrease in ARPE-19 cell metabolism after doxorubicin-HCl treatment. At 24 hours post-treatment, cells treated with 0.1 μg/mL and 10 μg/mL doxorubicin-HCl exhibited a significant decrease in metabolism (0.1 μg/mL, p=0.0246; 10 μg/mL, p<0.0001, Figure 3a). Interestingly, cells treated with 100 μg/mL doxorubicin-HCl demonstrated a significant increase in metabolism compared to the untreated control at 24 hours post-treatment (p=0.0002, Figure 3a). At 48 hours post-treatment, all doxorubicin-HCl-treated cells demonstrated significant decreases in metabolism compared to the control (0.01 μg/mL, p=0.0231; 0.1 μg/mL, p<0.0001; 1 μg/mL p<0.0001; 10 μg/mL, p<0.0001; 100 μg/mL, p<0.0001, Figure 3b). Of all the doxorubicin-HCl concentrations that were tested, 100 μg/mL led to the largest decrease in cell metabolism at the 48-hour time point (p<0.0001, Figure 3b).

**Figure 3:**
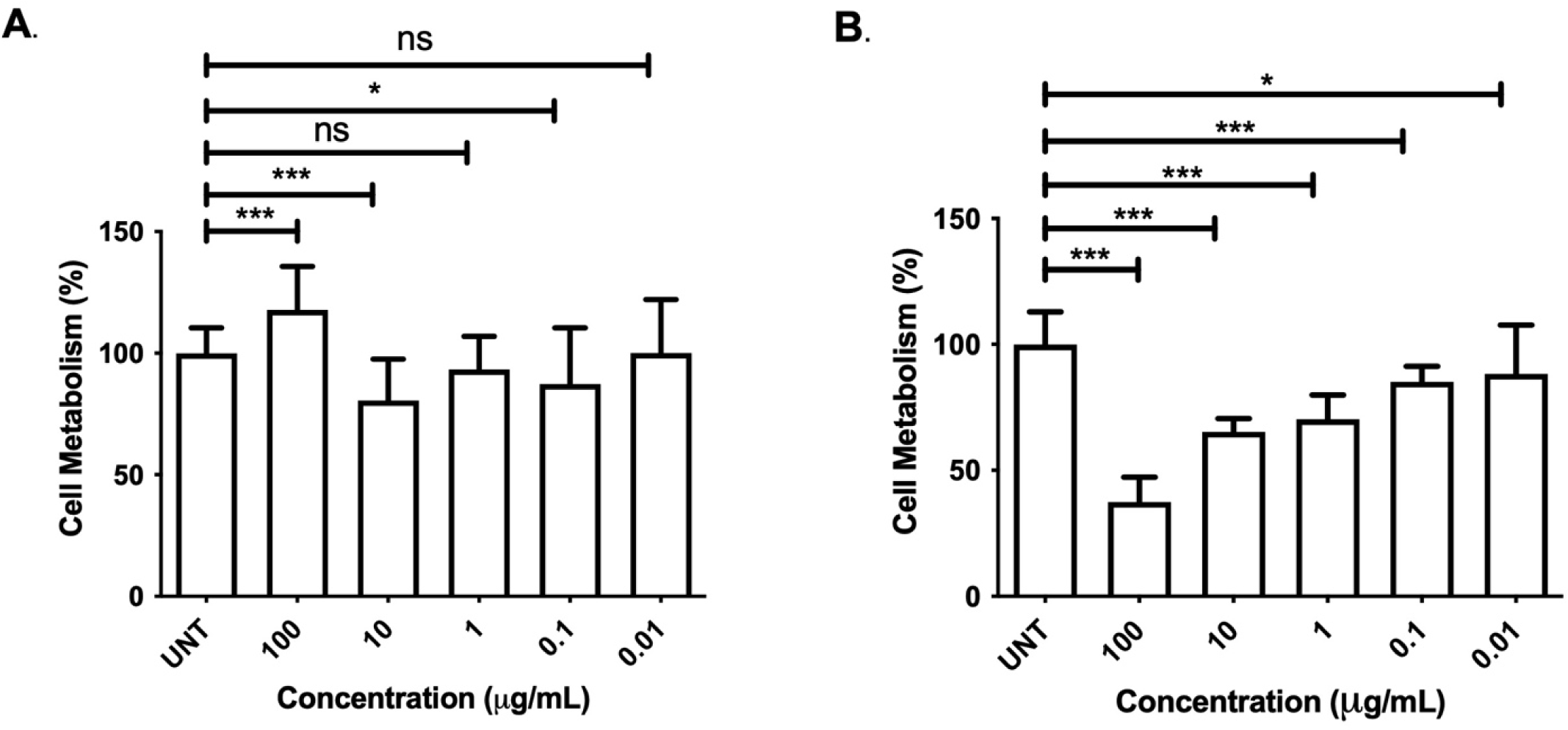
Doxorubicin-HCl toxicity in ARPE-19 cells. At 24 hours post-incubation (A), results were variable including a significant increase in cell metabolism at 100 μg/mL and decrease in metabolism at 10 μg/mL (100 μg/mL, p=0.0002; 10 μg/mL, p<0.0001). At 48 hours post-incubation (B), cells displayed a significant dose-dependent decrease in cell metabolism (0.01 μg/mL, p=0.0231; 0.1 μg/mL, p<0.0001; 1 μg/mL p<0.0001; 10 μg/mL, p<0.0001; 100 μg/mL, p<0.0001).

### Ultrasound- and Doxorubicin-HCl-treated Cells

ARPE-19 cells subject to both ultrasound and doxorubicin-HCl demonstrated greater decreases in cell metabolism compared to cells treated with doxorubicin-HCl alone. At 24 hours post-treatment, cells treated with both pulsed ultrasound and 0.1 μg/mL to 100 μg/mL doxorubicin-HCl exhibited significant dose-dependent decreases in metabolism compared to doxorubicin-treated cells alone (0.1 μg/mL, p=0.0142; 1 μg/mL, p<0.0001; 10 μg/mL, p<0.0001; 100 μg/mL, p<0.0001; Figure 4a). Cells subject to 100 μg/mL doxorubicin-HCl and ultrasound did not exhibit rebound increases in metabolism at 24 hours post-treatment. After 48 hours post-treatment, cells treated with both pulsed ultrasound and doxorubicin-HCl displayed significant reductions in cell metabolism compared to doxorubicin only-treated controls at concentrations greater than 0.1 μg/mL (0.1 μg/mL, p=0.0040, 1 μg/mL, p<0.0001; 10 μg/mL, p<0.0001; 100 μg/mL, p<0.0001; Figure 4b). These observed reductions in cell metabolism followed a dose-dependent trend.

**Figure 4:**
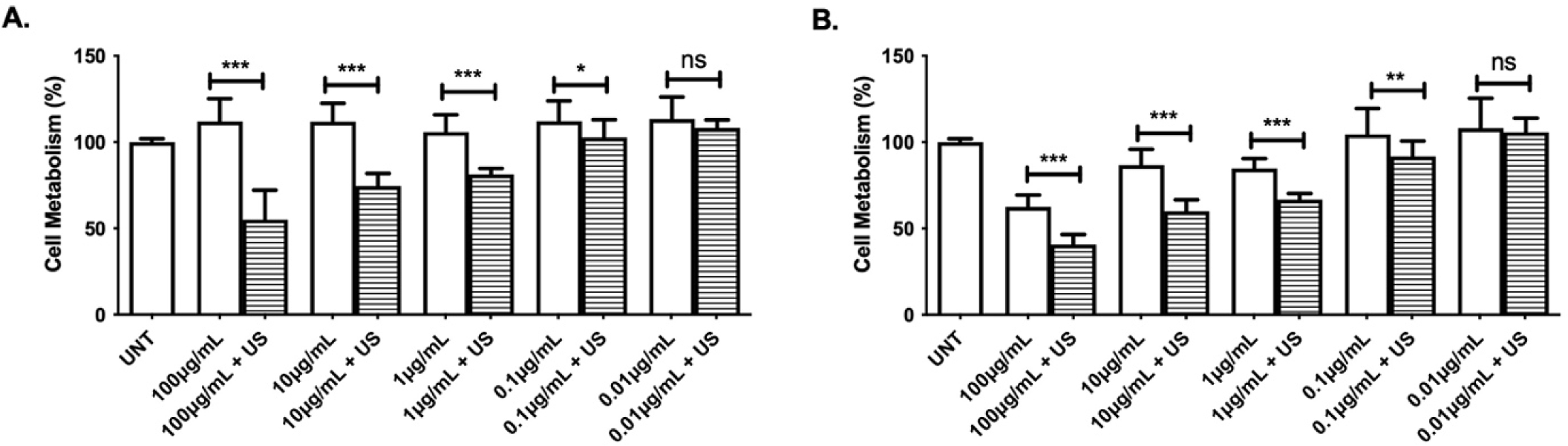
Ultrasound application with varying concentrations of doxorubicin-HCl at 24 hours (A) and 48 hours (B). The graphs demonstrate the effects of doxorubicin-HCl-treated cells with and without ultrasound exposure. When exposed to ultrasound, there was a significantly greater dose-dependent decrease in cell metabolism at 0.1 μg/mL or greater at both 24 and 48 hours (p<0.05). There was no effect visualized on 0.01 μg/mL of doxorubicin-HCl (24 hours, p=0.1488, 48 hours, p=0.607).

Trypan blue exclusion assay revealed no significant difference in cell viability between ARPE-19 cells treated with both 1 μg/mL doxorubicin-HCl and pulsed ultrasound and cells treated with 1 μg/mL doxorubicin-HCl only (24 hours, p=0.995; 48 hours, p=0.937; Figure 5a). A decrease in cell viability was observed in cells treated with 100 μg/mL doxorubicin-HCl and ultrasound compared to controls treated with doxorubicin-HCl only, but this difference was not significant (24 hours, p=0.606; 48 hours, p=0.794; Figure 5).

**Figure 5:**
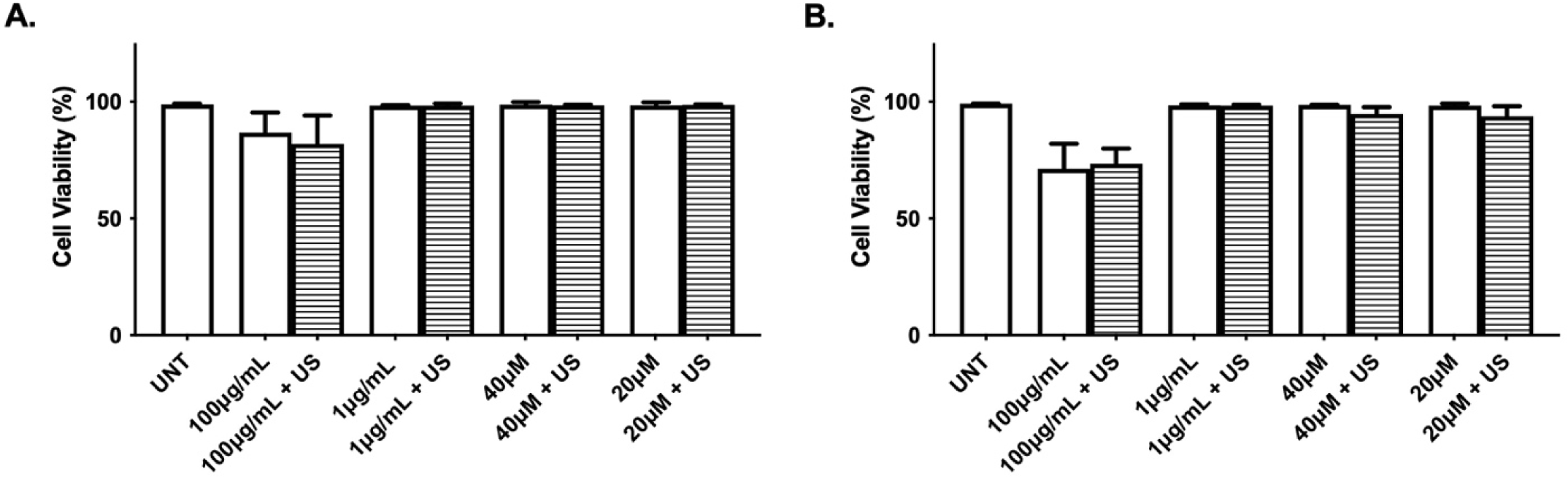
Trypan Blue analysis of doxorubicin- and cisplatin-treated cells with and without ultrasound (US) exposure at 24 hours (A) and 48 hours (B). 100 μg/mL and 1 μg/mL of doxorubicin-HCl and 40μM and 20 μM cisplatin were tested with and without pulsed ultrasound. At both 24 and 48 hours, there was no significant differences seen between cells treated with both doxorubicin-HCl and ultrasound and the doxorubicin-treated control (1 μg/mL: 24 hours, p=0.995; 48 hours, p=0.937; 100 μg/mL: 24 hours, p=0.606; 48 hours, p=0.794). There was no significant relationship seen between cisplatin-treated and ultrasound-treated cells and the cisplatin-treated controls (20 μM: 24 hours, p=0.870; 48 hours, p=0.151; 40 μM: 24 hours, p=0.715; 48 hours, p=0.0863).

### Ultrasound- and Cisplatin-treated Cells

At 24- and 48-hours post-cisplatin treatment, ARPE-19 cells demonstrated variable responses in cell metabolism with and without ultrasound exposure. At 24 hours post-treatment, there was no significant difference in metabolic activity between cisplatin and ultrasound-treated cells and control cells treated with cisplatin only (20 μM, p=0.511; 40 μM, p=0.722; Figure 6a). At 48 hours post-treatment, ultrasound-treated cells that were subsequently subject to 20 and 40 μM of cisplatin demonstrated statistically significant increases in metabolism compared to control cells treated with cisplatin only (20 μM, p<0.0001; 40 μM, p<0.0001, Figure 6b). To gain further insight into this apparently augmented metabolic response, we exposed our cisplatin solution to 1 MHz ultrasound at a PRF of 30 Hz in the absence of cells. Ultrasound-treated cisplatin (US-Cis) was then plated with ARPE-19 cells. There was no significant difference in metabolism between cells treated with 40 μM US-Cis and control cells treated with cisplatin only, while cells treated with 20 μM US-Cis displayed a significant decrease in metabolism compared to control cells treated with cisplatin only (40 μM US-Cis, p=,0.4004; 20 μM US-Cis, p=0.0093; Figure 6c). At 24- and 48-hours post-treatment, Trypan Blue assay revealed no significant difference in cell viability between cells treated with ultrasound plus cisplatin compared to control cells treated with cisplatin only (20 μM: 24 hours, p=0.870; 48 hours, p=0.151; 40 μM: 24 hours, p=0.715; 48 hours, p=0.0863; Figure 5).

**Figure 6:**
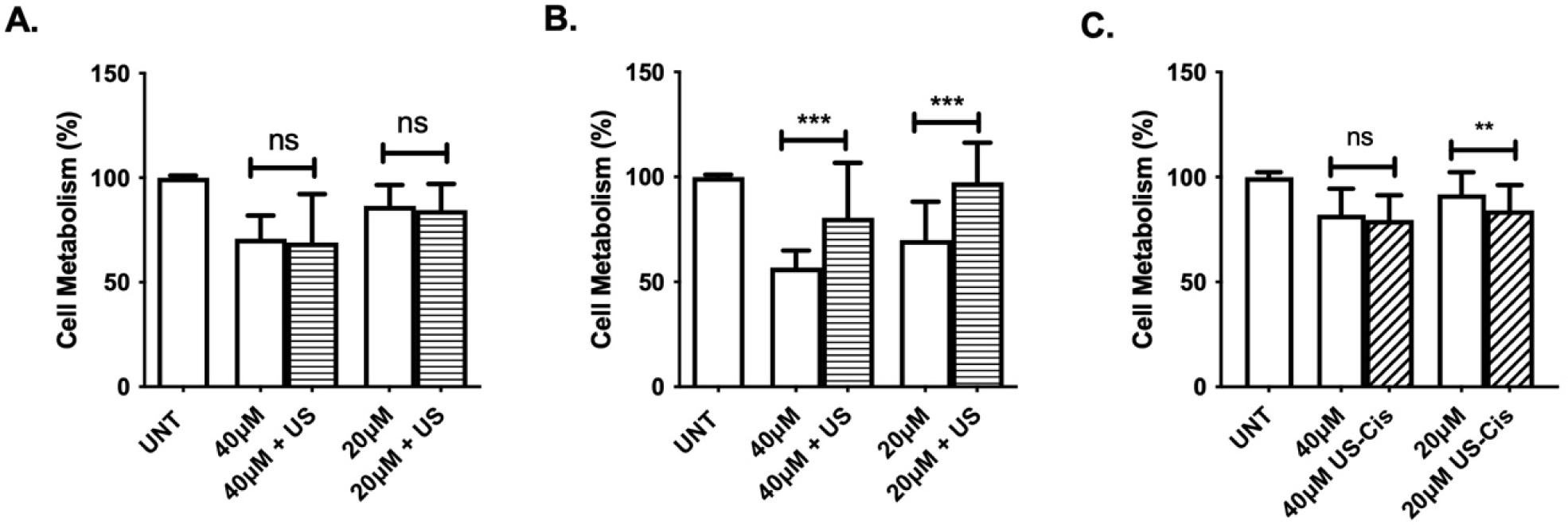
Effects of ultrasound plus cisplatin (A, B) on ARPE-19 cells at 24 hours (A), 48 hours (B). Panel C shows cells treated with US-Cis (cisplatin exposed to US prior to cell plating) after 24 hours. There were no significant effects seen at 24 hours post-incubation (A). After 48 hours, ultrasound-treated plus cisplatin-treated cells had increased cell metabolism compared to the metabolism of the cisplatin-treated controls (p<0.0001) (B). 20 μM concentrations of US-Cis had a decrease in cell metabolism (p=0.0093) (C). No significant difference in metabolism was found between 40 μM US-Cis and cisplatin-treated cells (p=0.4004, C).

## Discussion

The salient findings of the present investigation are: 1) dose-dependent effects of doxorubicin-HCl and cisplatin on ARPE-19 cell metabolism, 2) ultrasonic enhancement of doxorubicin-HCl toxicity, 3) ultrasonic modulation of cisplatin toxicity, 4) a nonsignificant effect of pulsed ultrasound on cisplatin- and doxorubicin-treated cells on cell death and 5) a viable pulsed ultrasound apparatus for use *in vitro*. These findings underscore the ability of ultrasound to regulate the cytotoxicity of antineoplastic agents, in a drug-specific manner, in cultured retinal epithelial cells and demonstrate its potential as a modulator of drug potency.

Because antineoplastic drugs such as doxorubicin-HCl and cisplatin are broadly used in clinical medicine, it is important to identify their secondary effects on tissues such as the eye. In our study, ARPE-19 cells demonstrated a dose-dependent decrease in metabolism after doxorubicin-HCl and cisplatin treatment alone, mimicking trends found in the literature [13,14]. Specifically, after a 48-hour incubation period, the dose-dependent decrease caused by doxorubicin-HCl mirrored previously published results of retinal epithelium cells treated with liposomal doxorubicin [13]. Interestingly, there was a deviation from this trend at the highest dose of doxorubicin-HCl; cells treated with 100 µg/mL doxorubicin-HCl exhibited a paradoxical increase in metabolic activity 24 hours post-treatment. Such findings were not seen in studies of NIH3T3 (mouse fibroblast) and MCF7 (breast adenocarcinoma) cell lines that were treated with 50 µg/mL doxorubicin for a 24-hour incubation period [17], and there are minimal to no published studies that identify such a trend. These data suggest that the cytotoxic effect of doxorubicin-HCl is dose-dependent at 48 hours but not at 24 hours post-treatment in ARPE-19 cells. This distinction may be caused by a unique activating response in cell metabolism upon treatment with an overwhelming dose of toxin; therefore, our observations may provide insight into how ARPE-19 cells respond to stress. It is also important to note that cells subject to the highest concentration of doxorubicin-HCl with ultrasound did not demonstrate the same paradoxical increase in metabolic activity seen at 24 hours post-treatment in the doxorubicin-HCl-only group. We suspect this response may have occurred prior to the 24-hour time point or failed to occur at all. Further studies are necessary to explore and validate these claims.

Cells treated with both ultrasound and doxorubicin-HCl displayed dose-dependent decreases in metabolic activity that were similar to those seen in cells treated with doxorubicin-HCl alone at 48 hours post-treatment. This toxicity was significantly amplified at both 24 and 48 hours after dual treatment, suggesting that ultrasound magnifies the cytotoxic effects of doxorubicin-HCl. These findings corroborate reports in the literature demonstrating that ultrasound enhances the potency of adriamycin on ovarian cancer *in vivo* [18]. Because ultrasound treatment amplified the magnitude of doxorubicin-HCl toxicity rather than the pattern of effect, it can be concluded that ultrasound did not change doxorubicin-HCl’s functional structure.

One possible explanation for the increased cytotoxicity of doxorubicin-HCl in the presence of ultrasound involves enhanced cellular exposure to doxorubicin-HCl via ultrasound-induced cell cavitation. Similar findings have been observed in previously published work, in which ultrasound was shown to enhance the corneal permeability of dexamethasone sodium *in vivo*, a phenomenon theorized to be mediated by stable and inertial cavitation [10]. This hypothesis of increased drug uptake has implications surrounding doxorubicin-HCl administration in several tissue types. Because cells subject to both ultrasound and doxorubicin-HCl exhibit a greater reduction in cell metabolism at all tested concentrations, we argue that a lower dose of doxorubicin-HCl could be combined with ultrasound to induce a cytotoxic effect comparable to that of a higher dose of doxorubicin-HCl alone. Lowering doxorubicin-HCl doses in conjunction with sonication to tissues of interest could minimize the adverse systemic side effects of doxorubicin-HCl while preserving its therapeutic effect.

In contrast to cells treated with doxorubicin-HCl, ARPE-19 cells treated with both cisplatin and ultrasound demonstrated a different response. Rather than amplifying cisplatin cytotoxicity and decreasing cell metabolism, ultrasound appeared to impart a cytoprotective effect. One theory regarding this increase in cell metabolism involves a physical modification of cisplatin’s molecular structure. To further explore this concept, we evaluated the ARPE-19 cell metabolic response to ultrasound-treated cisplatin (i.e., US-Cis). When cisplatin was pre-treated with ultrasound before exposure to ARPE-19 cells, the previously described protective effect was lost. There may be a component of this apparent ultrasound-induced abrogation of cisplatin toxicity that relies on the timing and duration of ultrasound exposure. Work done by a different group found that low-intensity ultrasound increases the apoptotic rate of cisplatin-treated hepatocellular carcinoma cells [18]. This finding suggests the type of sonication may also influence the cytotoxic effect of cisplatin. These findings also have important clinical implications. To minimize systemic side effects, ultrasound could conceivably be administered alongside cisplatin in ocular tissues that are particularly susceptible to cytotoxicity (e.g., retinal tissue) during cisplatin infusion.

It is important to note that ultrasound did not have a significant effect on doxorubicin- and cisplatin-treated ARPE-19 cell death. This suggests ultrasonic waves may prompt a nonfatal cellular response that transiently halts metabolism when exposed to these antineoplastic agents. Additional studies are necessary to further understand why sonication in the presence of doxorubicin-HCl and cisplatin appears to modulate cell metabolic activity but spares cell death.

Our study also demonstrates a viable ultrasound apparatus for the application of pulsed ultrasound *in vitro*. The observed decrease in ARPE-19 cell metabolism at 100 Hz ultrasound was suggestive of ultrasonic penetrance through the liquid-polystyrene-liquid interface, validating our pulsed ultrasound apparatus. Our sealed cell culture setup not only provides favorable parameters to test ultrasound penetration but also may have other applications in ocular drug delivery. Most notably, plate inversion allows for gas-filled particles, such as microbubbles, to rise to the cellular layer and facilitate optimal cell-particle contact. This minimizes the distance between drug and cells upon insonation.

Although there is growing interest in utilizing drug-filled microbubbles with insonation as a mechanism for targeted drug delivery, there remains a need to characterize the relationship between ultrasound and microbubble cargo in the absence of microbubbles [20]. Our study reveals ultrasound variably influences the potency of cisplatin and doxorubicin-HCl *in vitro*. Future microbubble experiments should explore the effects of ultrasound on the loading medication prior to involving microbubbles. Such information is crucial to establishing our understanding of the therapeutic role of ultrasound as well as augmenting ultrasound’s role as an ocular tool.

## Conclusion

Pulsed ultrasound enhances the cytotoxicity of doxorubicin-HCl at 24 and 48 hours post-treatment but abrogates cisplatin toxicity 48 hours post-treatment. This suggests ultrasound modulates cell-drug interactions in a drug-specific manner. These findings may influence the future development of ultrasound-assisted ocular drug delivery systems.

## Acknowledgments

This work was supported by the Fritch family endowment grant and Research to Prevent Blindness grant. The authors have no relevant financial or non-financial interests to disclose.

## Data Availability Statement

Data supporting this study are available upon request by contacting the corresponding author.

